# Immune chromatin reader SP140 regulates microbiota and risk for inflammatory bowel disease

**DOI:** 10.1101/2022.03.29.486273

**Authors:** Isabella Fraschilla, Hajera Amatullah, Raza-Ur Rahman, Kate L. Jeffrey

## Abstract

Inflammatory bowel disease (IBD) is driven by host genetics and environmental factors, including commensal microorganisms. Epigenetics facilitate integration of environmental cues for transcriptional output. However, evidence of epigenetic dysregulation directly causing host-commensal dysbiosis and IBD is lacking. Speckled Protein 140 (SP140) is an immune-restricted chromatin ‘reader’ with homology to Autoimmune Regulator (AIRE). SP140 mutations associate with three immune diseases: Crohn’s disease (CD), multiple sclerosis (MS) and chronic lymphocytic leukemia (CLL), but disease-causing mechanisms remain undefined. Here we identify a critical immune-intrinsic role for SP140 in preventing expansion of inflammatory Proteobacteria, including *Helicobacter* in mice and *Enterobacteriaceae* in humans. Mice harboring altered microbiota due to hematopoietic Sp140 deficiency exhibited severe colitis which was transmissible upon co-housing and ameliorated with antibiotics. SP140 was critical for calibration of macrophage microbicidal responses required for normal host-commensal crosstalk and elimination of invasive pathogens. Mutations within this epigenetic reader may constitute a predisposing event in human diseases provoked by the microbiome, such as IBD and MS.

## Introduction

Complex immune diseases, including inflammatory bowel disease (IBD) and multiple sclerosis (MS) are multifactorial diseases that develop as a result of gene-environment interactions (Graham and Xavier, 2020). Genome-wide association studies have identified over 240 genetic risk alleles that are associated with the two types of IBD: Ulcerative Colitis (UC) and Crohn’s disease (CD) (Jostins et al., 2012; Liu et al., 2015; Rivas et al., 2011). Many of these risk alleles overlap with loci that associate with MS (International Multiple Sclerosis Genetics et al., 2013) suggesting common disease initiating mechanisms. One shared CD and MS risk allele is SP140. SP140 is an immune-restricted member of the Speckled Protein (SP) epigenetic ‘reader’ family, consisting of SP100, SP110, SP140 and SP140L, which have high sequence homology with Autoimmune Regulator (AIRE) (Fraschilla and Jeffrey, 2020). Epigenetic readers are diverse proteins with specialized docking domains that ‘read’ covalent modifications primarily on histones to regulate transcription (Arrowsmith et al., 2012). To achieve this function, SPs all contain 3 ‘reader’ domains: 1) a SAND domain (named after the few SAND domain-containing proteins: SP100, AIRE, NucP41/P75, and DEAF) that interacts with DNA directly or through protein-protein interactions, 2) a plant homeodomain (PHD) that docks to histone methylation, and 3) a bromodomain that binds acetylated histones (Bienz, 2006; Bottomley et al., 2001; Filippakopoulos et al., 2012; Waterfield et al., 2014). Although, the PHD of SP140 may be atypical by facilitating SUMOylation of the bromodomain and associating with SETDB1, a histone methyltransferase of H3K9 that promotes gene silencing (Garcia-Dominguez et al., 2008; Ivanov et al., 2007; Peng and Wysocka, 2008; Zhang et al., 2016; Zucchelli et al., 2019). In addition, SP140 contains a caspase activation and recruitment domain (CARD) involved in multimerization and co-localization to ‘speckled’ promyelocytic leukemia (PML) nuclear bodies, macromolecular multiprotein complexes with diverse functionality including transcriptional repression (Corpet et al., 2020; Hoischen et al., 2018; Huoh et al., 2020).

Consistent with the predicted role for SP140 in gene silencing, in human macrophages, SP140 was found to predominantly occupy and maintain inaccessibility of promoters of developmentally silenced loci, such as the *HOXA* cluster (Mehta et al., 2017). *HOXA9* is a known promoter of stem-like state in hematopoietic stem cells (HSCs) and an inhibitor of macrophage differentiation and glycolysis (Argiropoulos and Humphries, 2007; Huang et al., 2012; Thorsteinsdottir et al., 2002; Zhou et al., 2018). *HOXA9* is therefore normally silenced in mature macrophages (De Santa et al., 2007), but not in SP140-deficient mouse or human macrophages, that ultimately display defective transcriptional responses to bacteria or viral ligands (Mehta et al., 2017). A global proteomics analysis subsequently found that SP140 directly represses topoisomerases to maintain heterochromatin, gene silencing, and macrophage responsiveness (Amatullah et al., 2021). Furthermore, an array of studies has now implicated SP140 as an essential factor in antibacterial, antiviral, and antiparasitic responses (Ji et al., 2021; Madani et al., 2002; Matsushita et al., 2021; Regad and Chelbi-Alix, 2001). However, despite some progress in understanding the function of this enigmatic epigenetic reader in macrophages, the role of SP140 in the immune response to intestinal microbiota, and how disruption of this function due to human genetic variation in SP140 may drive development of Crohn’s disease or MS remains unclear.

Development of complex immune disorders, such as IBD and MS, is dependent on both host genetics as well as cues derived from intestinal microbes and their metabolites (Amatullah and Jeffrey, 2020; Blander et al., 2017; Graham and Xavier, 2020; Iliev and Cadwell, 2021; Kaiko et al., 2016; Schulthess et al., 2019; Vinolo et al., 2011; Wu et al., 2020). Innate immune pathways are crucial for integrating bacterial, viral and fungal signals from the intestinal microenvironment to regulate gene expression, microbial balance and intestinal homeostasis (Adiliaghdam et al., 2021; Iliev and Cadwell, 2021; Schulthess et al., 2019). Moreover, many IBD-associated mutations are within genes essential for innate immune defense pathways, emphasizing their requirement for maintaining homeostatic host-microorganism relationships (Graham and Xavier, 2020; Jostins et al., 2012; Kugelberg, 2014). CD-associated mutations in *SP140* result in defective mRNA splicing and a reduction in SP140 protein (Matesanz et al., 2015; Mehta et al., 2017) rendering innate immune cells hypo-responsive. Further, mouse models support that SP140 is normally required for intestinal homeostasis via an innate immune function, as hematopoietic knockdown (Mehta et al., 2017) or whole mouse deletion of Sp140 exacerbates dextran sulfate sodium (DSS)-induced colitis in a manner dependent on macrophages (Amatullah et al., 2021). In this study, we investigated the possible link between intestinal microbial communities and SP140 control of intestinal homeostasis. We found that disruptions in SP140 result in a defective microbicidal transcriptome in phagocytes but enhanced activation of CD8+ T cells in colonic lamina propria and colitis. Remarkably, these intestinal malfunctions were dependent on the expansion of Proteobacteria, including *Helicobacter*, that was transmissible upon co-housing with wild-type mice and treated with antibiotics. This was further corroborated in CD patients bearing SP140 loss-of-function mutations that displayed significant elevation in Proteobacteria *Enterobacteriaceae*. Taken together, we have identified a key innate immune-intrinsic role for epigenetic reader SP140 in preventing intestinal inflammation by restricting the pathological expansion of a common member of the intestinal microbiota.

## Results and discussion

### SP140 is essential for antibacterial pathways in mice and human

SP140 is an essential host regulator of antibacterial defense to *Mycobacterium tuberculosis (Mtb)* (Ji et al., 2021; Pan et al., 2005; Pichugin et al., 2009) and gram negative bacteria (*Salmonella enterica* serovar Typhimurium, *Escherichia coli*, and *Citrobacter rodentium*) (Amatullah et al., 2021). Therefore, we sought to understand the mechanism by which SP140 contributes to antibacterial host defense and how SP140 deficiency may impact intestinal microbial dysbiosis and inflammation in CD patients bearing loss-of-function mutations. We examined transcriptional programs of human peripheral blood mononuclear cells (PBMCs) obtained from healthy controls expressing wild-type SP140 (HC SP140^wt^), Crohn’s disease (CD) patients expressing wildtype *SP140* (CD SP140^wt^), or CD patients that were homozygous for CD-associated SP140 mutations (CD SP140^mut^), obtained through the Prospective Registry in IBD Study at MGH (PRISM) cohort. A prominent gene set downregulated in CD SP140^mut^ cells compared to both healthy controls and CD *SP140^wt^* PBMCs included genes associated with bacterial defense (**Fig. 1A**). CD SP140^mut^ cells had reduced expression of transcripts encoding for subunits of NADPH oxidase (*CYBB, NCF1, NCF2, NCF4*) and H^+^-ATPase (*ATP6V0A1, ATP6V0D1*), which are known complexes that act at the phagosome to produce antibacterial reactive oxygen species (ROS) and low pH, respectively. Moreover, at steady state, genes essential for the degradation or killing of bacteria, including calprotectin subunits (*S100A8, S100A9*), lysozyme (*LYZ*), and cathepsins (*CTSB, CTSH*), were also reduced in CD SP140^mut^ cells. Since calprotectin is a well-studied antimicrobial protein capable of enhancing bacterial killing, we confirmed reduced protein levels of calprotectin component S100A8 in CD SP140^mut^ PBMCs compared to SP140^wt^ CD patient donors (**Fig. 1B**).

**Figure 1.**
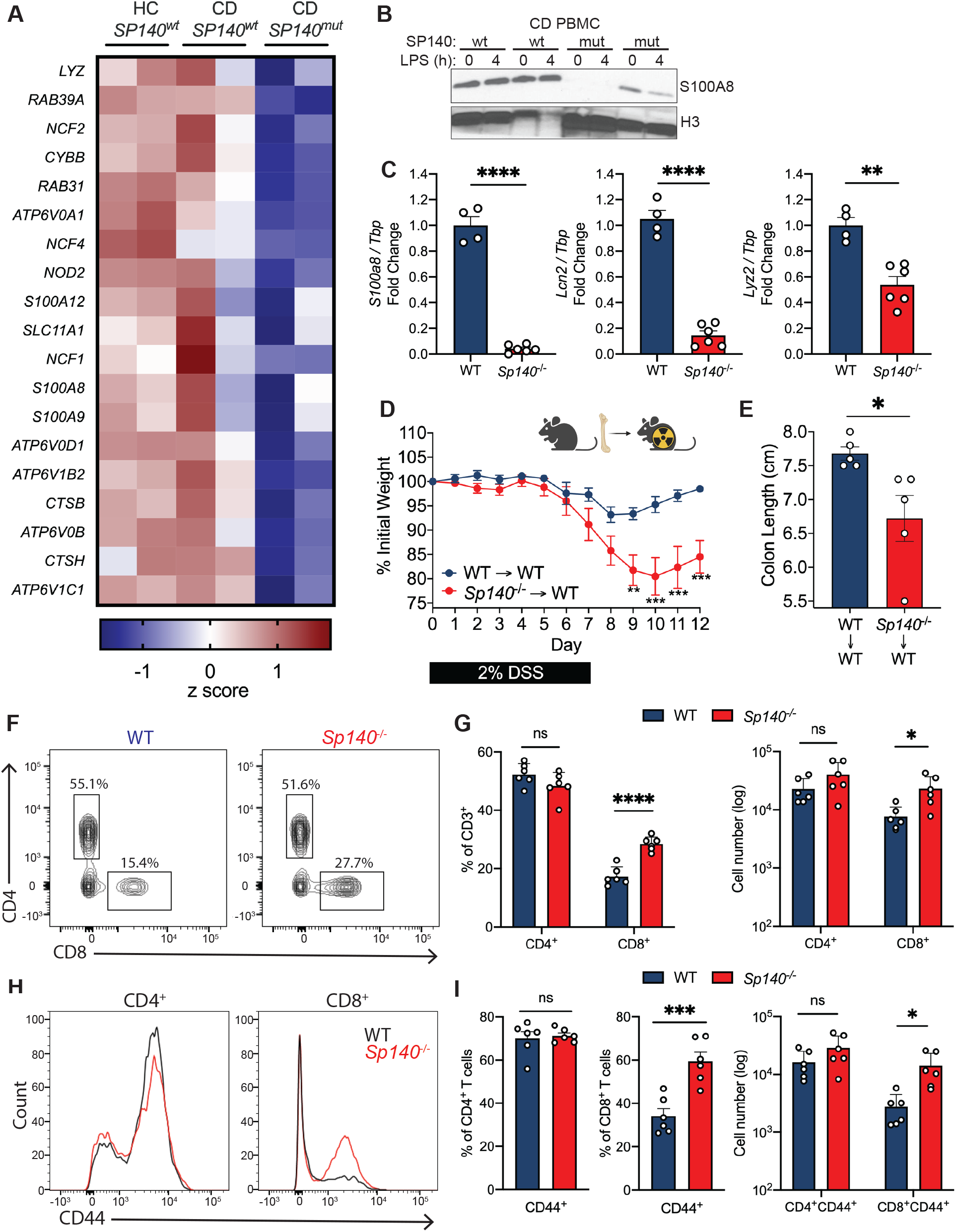
SP140 is essential for antimicrobial gene programs in mouse and human. **(A)** Heat map of genes down-regulated (log_2_FC>2) in human peripheral blood mononuclear cells (PBMC) from healthy controls (HC), Crohn’s disease (CD) patients bearing wildtype *SP140* (*SP140^wt^*) or CD-risk *SP140* genetic variant (*SP140^mut^*) (n=2, RNA-seq) from <u>GSE89876</u>. **(B)** Immunoblot of S100A8 in PBMC from *SP140^wt^* or *SP140^mut^* CD patients after 0 or 4 hours LPS stimulation (100 ng/mL). **(C)** Expression of mouse *S100a8, Lcn2*, and *Lyz2* mRNA relative to *Tbp*, as determined by qPCR, in mouse bone marrow-derived macrophages (BMDMs) from wildtype (WT) and *Sp140*^-/-^ mice. **(D)** Daily body weight of WT and *Sp140*^-/-^ mice and **(E)** quantification of day 12 colon lengths after 2% dextran sodium sulfate (DSS) administration (n=5). **(F-I)** Flow cytometry analysis of WT and *Sp140*^-/-^ mouse colon lamina propria (n=6). **(F)** Representative flow cytometry plots of live CD45^+^CD3^+^ cells gated on CD4^+^ and CD8^+^ T cells. **(G)** Quantification of frequency and total count of CD4^+^ and CD8^+^ T cells. **(H)** Representative histogram overlay of CD44 expression in CD4^+^ and CD8^+^ T cells. **(I)** Quantification of frequency and total count of CD44^+^CD4^+^ and CD8^+^ T cells. Data are mean of 2-6 biological replicates. *P<0.05, **P<0.01, ***P< 0.001, ****P<0.0001; unpaired *t* tests for (C), (E), (F), (H), two-way ANOVA for (D). Error bars represent s.e.m.

Human and mouse SP140 are only 54% homologous at the amino acid level, owing to inclusion of an intrinsically disorder region (IDR) in human SP140 (Fraschilla and Jeffrey, 2020). However, we found that antimicrobial transcripts, such as *S100a8, Lyz2* and *Lcn2*, were also downregulated in *Sp140*^-/-^ bone-marrow derived macrophages (BMDMs) (**Fig. 1C**). The conserved role of SP140 in controlling antimicrobial gene programs suggests that this chromatin reader plays an essential role in host defense. Since previous analysis of genome-wide occupancy of SP140 in human macrophages found that SP140 did not directly occupy antimicrobial gene loci and promoted gene repression at silenced chromatin regions (Mehta et al., 2017), SP140 likely promotes an antibacterial gene program via maintenance of heterochromatin and macrophage identity.

SP140 expression has been shown to be immune-restricted and particularly high in macrophages (Mehta et al., 2017). Intestinal macrophages, which differentiate from circulating monocytes (Bain et al., 2014), and other phagocytes in the intestine are responsible for bactericidal responses, namely production of ROS and antimicrobial proteins (Schulthess et al., 2019; Smythies et al., 2005). Furthermore, other IBD-associated loci have similarly been shown to regulate microbial responses in the intestine via epithelial cells (Conway et al., 2013; Graham and Xavier, 2020; Ramanan et al., 2014), which are resistant to irradiation. To determine whether SP140 expression in immune cells is sufficient to protect against intestinal inflammation, we irradiated wildtype (WT) mice and reconstituted them with WT or *Sp140*^-/-^ bone marrow. Mice with specific Sp140 deficiency in the hematopoietic compartment displayed significantly exacerbated DSS-colitis compared to WT controls, as determined by weight loss (**Fig. 1D**) and colon length (**Fig. 1E**). Thus, Sp140 within the immune compartment is essential for intestinal homeostasis. To extend our *ex vivo* characterization of antibacterial programs in mouse and human primary cells, we next investigated whether Sp140 deficiency alters the development of intestinal phagocyte populations that are necessary for bacterial clearance. We performed immunophenotyping of colonic lamina propria CD45^+^ cells by flow cytometry analysis in Sp140^-/-^ and WT mice. Sp140 deficiency did not affect the proportions of mononuclear phagocyte populations (**Fig. S1C**) or B cells (**Fig. S1A, B**) suggesting that recruitment and development of these cell types is intact in *Sp140*^-/-^ mice. However, a dysfunctional response to the microbiota can lead to adaptive immune cell infiltration in the gut (Feng et al., 2010). Indeed, we observed a significant and specific expansion of total CD8^+^ T cells (**Fig. 1F, G**) as well as a significant increase in the frequency and number of activated CD8^+^CD44^+^ T cells (**Fig. 1H, I**) in the colonic lamina propria of *Sp140*^-/-^ mice. Intriguingly, this defect was specific to CD8^+^ T cells as CD4^+^ T cells and CD4^+^CD44^+^ T cell frequencies and numbers were unaltered with Sp140 deficiency (**Fig. 1F-I**). Taken together, these results show that Sp140 is required for macrophage antibacterial responses and this defect leads to CD8^+^ T cell activation in the gut lamina propria prior to the manifestation of disease.

### Cohousing wild-type mice with *Sp140*^-/-^ mice increased severity of induced colitis

Mice with an shRNA-mediated hematopoietic knockdown of *Sp140* and *Sp140*^-/-^ mice generated by CRISPR/Cas9 targeting are more susceptible to DSS-mediated colitis, in a manner dependent on Sp140 control of cytokine production and bacterial killing in macrophages (Amatullah et al., 2021; Mehta et al., 2017). Further, an abundance of studies has demonstrated that mice deficient in innate immune system components, or cytokines, have altered microbiota with increased inflammatory capacity (Caruso et al., 2019; Elinav et al., 2011; Frederic et al., 2012; Ramanan et al., 2014; Zenewicz et al., 2013). Moreover, presence of activated CD8^+^ T cells in the colon is frequently the result of microbiota dysbiosis (Ramanan et al., 2014; Tanoue et al., 2019; Yu et al., 2020). We therefore questioned whether the microbiota of *Sp140*^-/-^ mice could contribute to the exacerbated colitis phenotype. We designed an experiment in which WT mice were cohoused with *Sp140*^-/-^ mice or were housed separately. After 4 weeks of cohousing, a period previously established to be sufficient for the transmission of microbiota between mice via coprophagia (Stappenbeck and Virgin, 2016; Zenewicz et al., 2013), we administered DSS and examined the resulting weight loss and inflammation associated with colitis. Whereas *Sp140*^-/-^ mice exhibit exacerbated colitis compared to WT mice when housed separately, cohousing WT mice with *Sp140*^-/-^ mice led to enhanced weight loss of WT mice, similar to separated or cohoused *Sp140*^-/-^ mice (**Fig. 2A, B**). To examine inflammation, we measured colon lengths at day 12 post-DSS administration and found significantly shorter colons in WT mice that were cohoused with the *Sp140*^-/-^ mice than WT mice that were housed separately (**Fig. 2C**). Histological examination of colons confirmed an increase in inflammation in WT mice specifically upon co-housing as shown by crypt elongation, thickening of the mucosa, and a leukocyte infiltrate to the level of the lamina submucosa (**Fig. 2D**). The transmission of the altered gut microbiota from *Sp140*^-/-^ mice to cohoused WT mice, along with increased susceptibility to DSS-induced colitis, indicates that the altered gut microbiota works as a contributing factor for, rather than a consequence of, the disease. To understand whether the microbiome of *Sp140*^-/-^ mice is colitogenic due to increased inflammatory capacity, we collected and filtered stool homogenates from untreated *Sp140*^-/-^ mice and controls and stimulated BMDMs *in vitro*. Stool homogenates obtained from *Sp140*^-/-^ mice proved to be hyperinflammatory compared to WT stool, as shown by a significant increase in the production of the pro-inflammatory cytokine Interleukin (IL)-6 (**Fig. 2E**).

**Figure 2.**
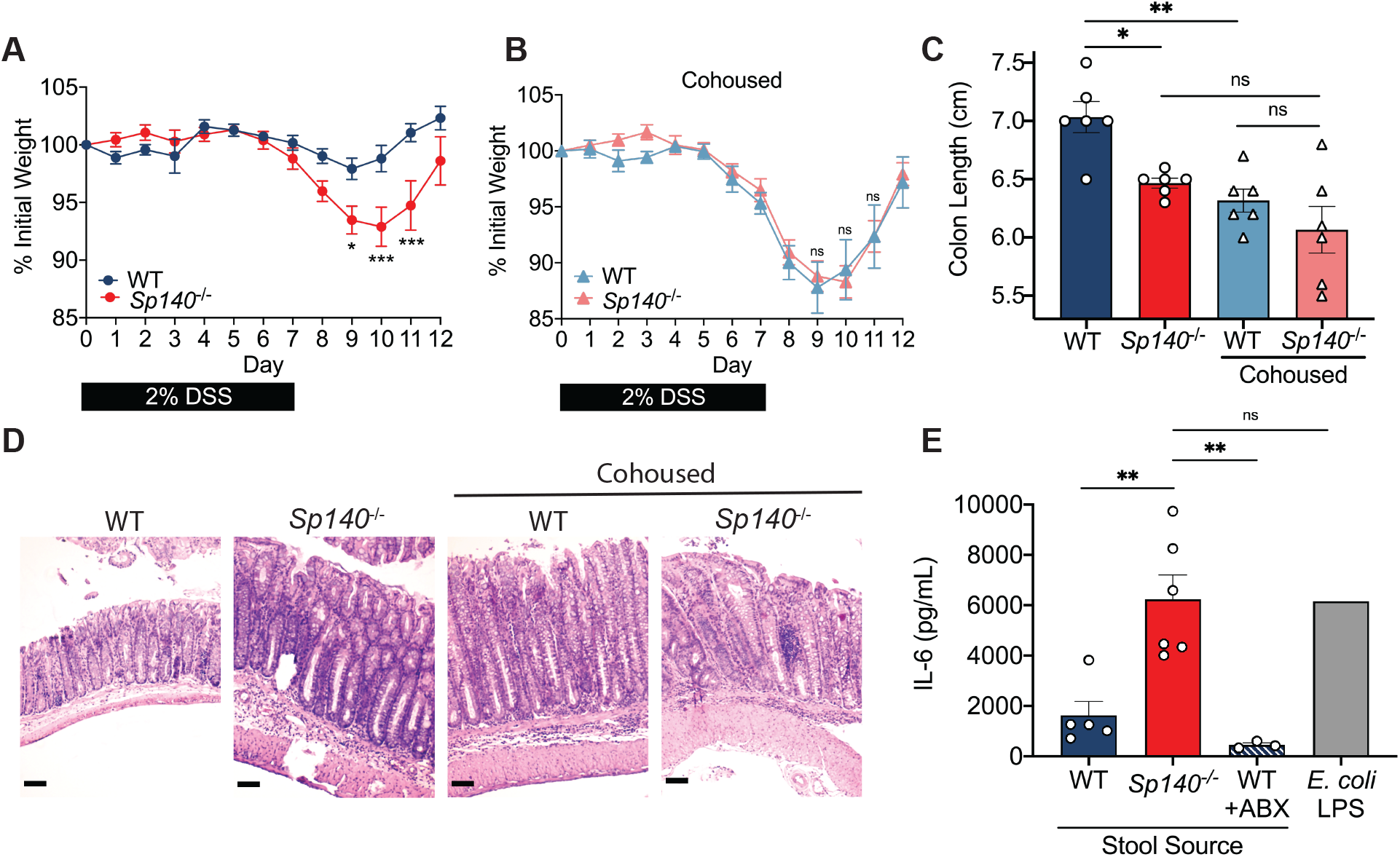
The pro-inflammatory intestinal microbiome of *Sp140*^-/-^ mice is transferable and exacerbates colitis. **(A)** Daily body weight of separately housed wildtype (WT) and *Sp140*^-/-^ mice after 2% dextran sodium sulfate (DSS) administration (n=6). **(B)** Daily body weight of cohoused WT and *Sp140*^-/-^ mice after 2% dextran sodium sulfate (DSS) administration (n=6). **(C)** Day 12 colon lengths of separated or cohoused WT and *Sp140*^-/-^ mice after 2% DSS administration (n=6). **(D)** Representative hematoxylin and eosin-stained sections of day 12 distal colon tissue from separated or cohoused WT and *Sp140*^-/-^ mice after 2% DSS administration. **(E)** IL-6 production, as determined by ELISA, of bone marrow-derived macrophages (BMDMs) supernatants after 16 hours stimulation with LPS (1mg/mL) or stool homogenates (1mg/mL) from WT (n=5), *Sp140*^-/-^ (n=6), or broad-spectrum antibiotic treated WT mice (n=3); representative of 3 experiments. *P < 0.05, **P < 0.01, ***P < 0.001, ****P<0.0001, two-way ANOVA for (A) and (B), one-way ANOVA for (C) and (E). Error bars represent s.e.m.

### *Sp140*^-/-^ mice have altered commensal microbiota that is transmissible

Our data support the hypothesis that *Sp140*^-/-^ mice have an altered microbiota that is transmissible to wild-type mice. Thus, we undertook bacterial 16S rRNA gene pyrosequencing to examine the microbiome in four groups of mice: WT mice, *Sp140*^-/-^ mice, and WT mice or *Sp140*^-/-^ mice cohoused with each other. Prior to DSS treatment we collected fecal samples from each mouse, prepared fecal DNA, and used barcoded pyrosequencing of the 16S rRNA gene V4 region to explore the change in the microbiome. A significant increase in community richness in *Sp140*^-/-^ mice was observed. Furthermore, this increase in richness was transferred to cohoused WT mice compared to non-cohoused WT controls (**Fig. 3A**). UniFrac-based principal component analysis (PCA) determined that WT and *Sp140*^-/-^ microbiome communities were significantly dissimilar from each other (**Fig. 3B**). Moreover, this difference was lost upon cohousing whereby WT microbiomes now overlapped with *Sp140*^-/-^ communities, demonstrating a dominance of *Sp140*^-/-^ microbiota (**Fig. 3B**). Proportional taxonomy analysis revealed that *Sp140*^-/-^ mouse stool contained a higher percentage of Proteobacteria phyla than WT mice, and Proteobacteria then bloomed in WT cohoused mice (**Fig. 3C**) thus identifying a putative, transmissible bacteria of the *Sp140*^-/-^ mouse microbiome that may confer sensitivity to colitis. Proteobacteria expansion is associated with worsened colitis in mouse models (Garrett et al., 2010; Ni et al., 2017). Furthermore, we specifically determined that *Sp140*^-/-^ intestines were colonized by a greater percentage of classes *Epsilonproteobacteria* and *Alphaproteobacteria* (**Fig. 3D**). Conversely, *Sp140*^-/-^ exhibited reduced Verrucomicrobia phyla that are present in wild-type controls (**Fig. 3C**). Unbiased Linear discriminant analysis Effect Size (LEfSe) analysis determined a significant elevation of colitis-driving *Helicobacter* genera of the *Epsilonproteobacteria* class and *Mucispirillum* genera of the Deferribacteria phylum in *Sp140*^-/-^ mice compared to WT controls (**Fig. 3E**). Notably, a significant reduction in the protective *Akkermansia muciniphila* was also observed in *Sp140*^-/-^ mice (**Fig. 3E**). Furthermore, the Sp140 deficient microbiome was efficiently transferred to cohoused WT mice (**Fig. 3F**) to the point where no significant differences were detected between *Sp140*^-/-^ mice and WT cohoused mice by LEfSe analysis (**Fig. 3G-H**). Thus, Sp140 deficiency triggers the development of a colitogenic microbiome, that is efficiently transferred upon cohousing.

**Figure 3.**
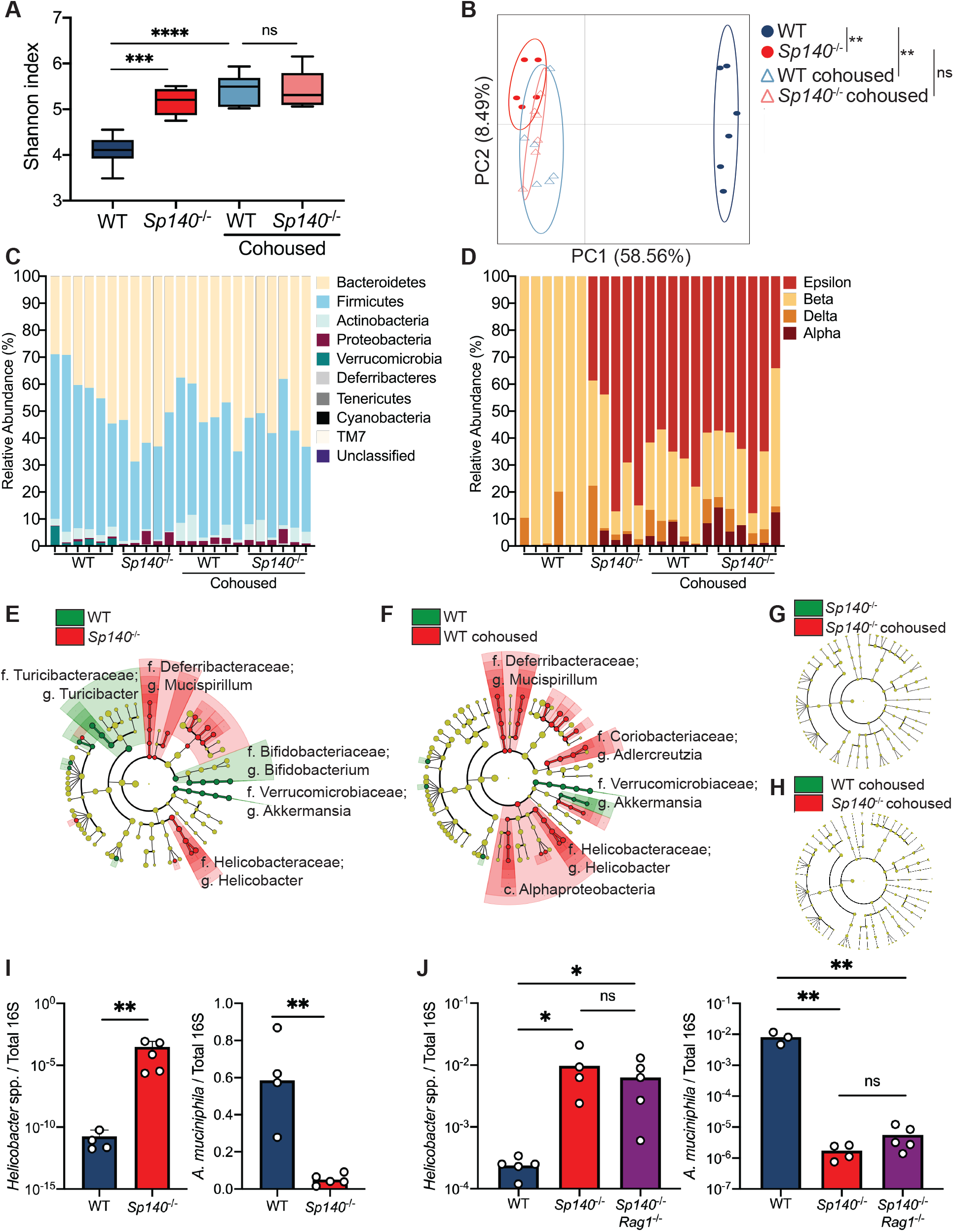
Sp140 deficiency in mice permits a *Helicobacter* bloom and a reduction in *Akkermansia*. **(A)** Diversity of fecal bacterial communities of separately housed or cohoused wildtype (WT) and *Sp140*^-/-^ mice as determined by Shannon Index. **(B)** Principal Coordinate Analysis (PCA), P<0.05, PERMANOVA on unweighted UniFrac distance. **(C)** Distribution of bacterial phyla operational taxonomic units presented as relative abundance in each sample. **(D)** Distribution of bacterial class operational taxonomic units presented as relative abundance of Proteobacteria in each sample. **(E)** The compositional differences between separately housed WT and *Sp140*^-/-^ mice, **(F)** separately housed WT and WT cohoused with *Sp140*^-/-^ mice, **(G)** separately house *Sp140*^-/-^ and cohoused *Sp140*^-/-^ mice, and **(H)** cohoused WT and *Sp140*^-/-^ mice were determined by a linear discriminant analysis using LEfSe (https://huttenhower.sph.harvard.edu/galaxy/). **(I)** Expression of *Helicobacter* species (spp.) and *Akkermansia muciniphila* 16S rRNA relative to total 16S rRNA, as determined by qPCR, in stool from WT and *Sp140*^-/-^ mice (n=4-5). **(J)** Expression of *Helicobacter* species (spp.) and *Akkermansia muciniphila* 16S rRNA relative to total 16S rRNA in stool from WT, *Sp140*^-/-^, and *Sp140^-/-^Rag1*^-/-^ mice (n=3-5). *P < 0.05, **P < 0.01, ***P < 0.001, ****P<0.0001. Error bars represent s.e.m. (I) unpaired *t* tests. (A, J) one-way ANOVA. *P< 0.05, **P<0.01, ***P<0.001, ****P<0.0001. Error bars represent s.e.m.

The *Helicobacter* genus can drive colitis and inflammation in mice (Cahill et al., 1997; Chai et al., 2017; Kullberg et al., 1998; Yang et al., 2013) while *A. muciniphila* promotes mucosal wound healing and ameliorates colitis in mouse models (Alam et al., 2016; Ansaldo et al., 2019). We therefore confirmed the significant elevation in *Helicobacter* and reduction in *A. muciniphila* in *Sp140*^-/-^ feces by qPCR (**Fig. 3I**). To assess if Sp140 control of microbiota was intrinsic to innate immune cells, or if there was a role for lymphocytes, we crossed *Sp140*^-/-^ mice with lymphocyte-deficient *Rag1*^-/-^ mice and found that feces from double mutant *Sp140^-/-^ Rag1*^-/-^ mice still displayed significantly elevated *Helicobacter* and reduced *A. muciniphila* (**Fig. 3J**). Thus, Sp140 deficiency specifically in innate cells drives proteobacteria expansion. Furthermore, this data demonstrates that the elevation in activated CD8^+^ T cells in the colonic lamina propria of *Sp140*^-/-^ mice (**Fig. 1F-I**) is indeed a consequence of microbiota disruption.

### Crohn’s disease patients with *SP140* mutations exhibit increase in intestinal *Enterobacteriaceae*

Our analysis of intestinal bacterial populations in *Sp140*^-/-^ mice suggest that a bloom of pathogenic Proteobacteria results in exacerbated colitis. To demonstrate that expanded Proteobacteria was a driving factor of exacerbated colitis in *Sp140*^-/-^ mice, we administered mice the antibiotic metronidazole via drinking water for 2 weeks before DSS-colitis induction. Indeed, metronidazole treatment protected *Sp140*^-/-^ mice from inflammation associated with colitis as determined by increased colon length (**Fig. 4A**) and reduced crypt hyperplasia (**Fig. 4B**) compared to *Sp140*^-/-^ mice that did not receive antibiotics. Metronidazole targets gram-negative anaerobes, including species within the Proteobacteria phylum, and is used in the clinic to treat CD. In order to determine whether manipulation of the microbiota might be a beneficial route of treatment in CD patients bearing *SP140* loss-of-function mutations, we sought to determine whether the architecture of microbial communities in *Sp140*^-/-^ mice recapitulated species dominance in humans with SP140 loss-of-function. We performed 16S sequencing analysis of stool obtained from healthy controls, CD SP140^wt^, and CD SP140^mut^ individuals (**Fig. S2A**). Alpha-diversity analysis determined that stool from CD patients exhibited reduced Shannon diversity compared to healthy controls (**Fig. 4C**) as previously reported (Franzosa et al., 2019). However, each group segregated from each other in principal component analysis (**Fig. 4D**) demonstrating that while both CD SP140^wt^ and CD SP140^mut^ individuals lose bacterial diversity, there are compositional differences between these two groups of CD patients. Proportional taxonomy analysis demonstrated that Proteobacteria diversity was lost in CD SP140^mut^ individuals whereby Gammaproteobacteria dominates (**Fig. 4E**). Specifically, we observed a significant increase in *Enterobacteriaceae*, a family of Gammaproteobacteria, in CD patients bearing SP140 loss-of-function mutations (**Fig. 4F**). The *Enterobacteriaceae* family of the Proteobacteria phylum encompasses gram negative facultative anaerobes including *Escherichia coli, Citrobacter rodentium*, and *Salmonella enterica*. Blooming of facultative anaerobes, particularly *Enterobacteriaceae*, with a concomitant depletion of obligate anaerobes associated with short chain fatty acid production is a common feature of IBD gut microbiomes (Knights et al., 2014; Lloyd-Price et al., 2019; Morgan et al., 2012). Furthermore, *Enterobacteriaceae* blooms are associated with chronic inflammation and treatment failure in IBD (Olbjørn et al., 2019; Walujkar et al., 2014). We predict that the inability to initiate production of toxic reactive oxygen species and antibacterial proteins in SP140-deficient CD patients permits expansion of *Enterobacteriaceae* as has been suggested for individuals with loss-of-function mutations in NAPDH oxidase subunits (Denson et al., 2018; Muise et al., 2012; Plichta et al., 2019).

**Figure 4.**
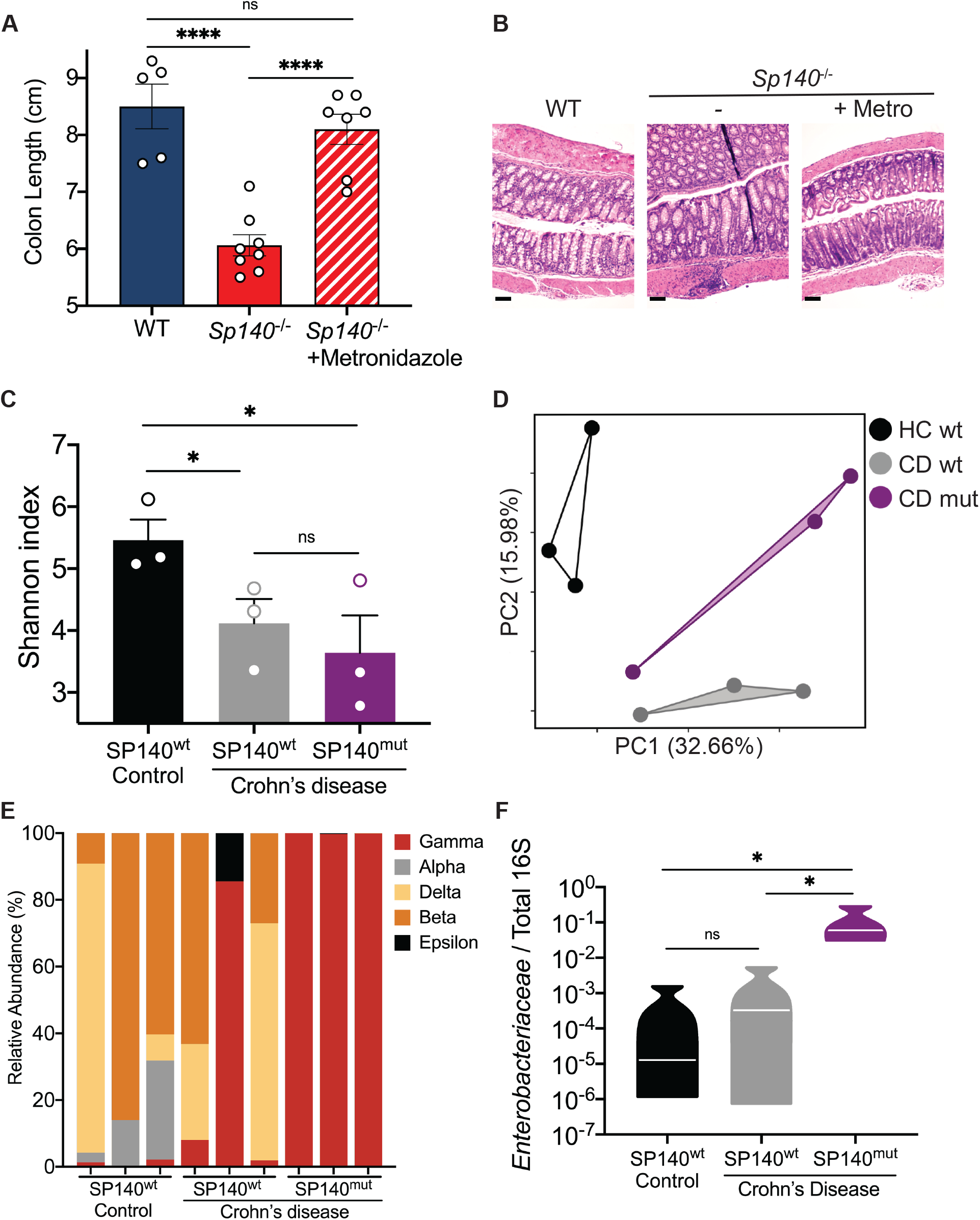
CD-associated SP140 loss-of-function variant associates with increased intestinal *Enterobacteriaceae* in humans. **(A)** Day 12 colon lengths of WT, *Sp140*^-/-^, and *Sp140*^-/-^ mice treated with or without metronidazole and administered 2% dextran sodium sulfate (DSS). **(B)** Representative hematoxylin and eosin-stained sections of day 12 distal colon tissue after 2% DSS administration to WT, *Sp140*^-/-^, and *Sp140*^-/-^ mice treated with metronidazole (metro). **(C)** Diversity of fecal bacterial communities of healthy controls, Crohn’s disease (CD) patients expressing wildtype SP140 (SP140^wt^), and CD patients bearing the common SP140 genetic variant (SP140^mut^). **(D)** Principal Coordinate Analysis (PCA), P<0.05, PERMANOVA on unweighted UniFrac distance. **(E)** Distribution of bacterial class operational taxonomic units presented as relative abundance of Proteobacteria in each sample. **(F)** Expression of *Enterobacteriaceae* 16S rRNA relative to total 16S rRNA in human stool. (A-B) n=5-8; (C-F) n=3, one-way ANOVA. *P < 0.05, **P < 0.01, ***P < 0.001, ****P<0.0001. Error bars represent s.e.m.

IBD symptoms and severity are a spectrum, suggesting that subtypes of disease are dependent on an axis of genetics, environmental cues, and epigenetics. Understanding how risk alleles contribute to pathogenicity and how epigenetic dysregulation is a contributing factor to IBD is essential for identifying disease subtypes as well as developing precision medicine treatment options. Here we have identified a key regulatory role for a novel epigenetic reader protein, SP140, that is also a risk allele for IBD, in the maintenance of macrophage function and prevention of intestinal microbe expansion, specifically Proteobacteria members. These findings have important implications for not only CD patients with loss-of-function SP140 mutations but also for MS that is driven by an altered microbiota. Genetic loss of SP140 has drastic consequences on the composition of the microbial communities leading to a shift towards a pro-inflammatory configuration that could predispose, and contribute to phenotypes of, both CD and MS (Berer et al., 2011) (Berer et al., 2017; Britton et al., 2019; Britton et al., 2020; Cekanaviciute et al., 2017), as well as other diseases influenced by genetics and microbe-derived factors. These results shed light on the etiology of CD and provide insight into the relationship between genetic susceptibility, epigenetics, microbial dysbiosis and the immune response in the gut. Patients with SP140 mutations may benefit from targeting pro-inflammatory commensals through targeted antibiotics or fecal transplants.

## Author Contributions

I.F. performed and interpreted *in vitro* and *in vivo* experiments, with help from H.A. R-U.R performed analysis of 16S sequencing. K.L.J conceived and supervised the study, acquired funding and wrote the final manuscript, along with I.F.

## Acknowledgements

We sincerely thank the clinical coordinators and patients enrolled in the Prospective Registry in IBD study at Massachusetts General Hospital (PRISM), Russell Vance (University of California, Berkeley) for *Sp140*^-/-^ mice, Amy Avery (Massachusetts General Hospital) for technical assistance with 16S sequencing, the entire Jeffrey lab for valuable scientific discussions as well as Robert Anthony, James Moon, Charles Evavold, and Jonathan Kagan for critical reading of the manuscript. This study was supported by F31DK127518 (I.F.), the Kenneth Rainin Foundation (Innovator and Synergy Awards to K.L.J), NIH R01DK119996 (K.L.J), K.L.J is a John Lawrence MGH Research Scholar 2020-2025.

## Materials and Methods

### Mice

C57BL/6J mice were originally purchased from Jackson Laboratory and bred in-house. All mice were bred and housed under specific pathogen-free conditions according to the National Institutes of Health (NIH). All animal experiments were conducted under protocols approved by the MGH Institutional Animal Care and Use Committee (IACUC), and in compliance and appropriate ethical regulations. For all experiments, age- and sex-matched mice were used. *Sp140*^-/-^ mice were made on C57BL/6J background as previously characterized (Ji et al., 2021). *Rag1*^-/-^ mice were purchased from Jackson Laboratory and crossed to *Sp140*^-/-^ mice. For cohousing experiments, WT and *Sp140*^-/-^ mice were cohoused from weaning for 4 weeks at a 2:2 ratio per cage.

### Human blood and stool samples

All human samples were collected under Institutional Review Board (IRB)-approved protocols by Massachusetts General Hospital (MGH), including informed consent obtained in accordance with relevant ethical regulations. Blood and stool samples were collected from patients that were enrolled in the “Prospective Registry in IBD Study at Massachusetts General Hospital (PRISM, IRB# FWA00003136). Study research coordinators obtained consent, and medical history was obtained and confirmed by review of the electronic medical record. Human peripheral blood mononuclear cells (PBMCs) were obtained from healthy human volunteers (Blood Components Lab, Massachusetts General Hospital) or from Crohn’s disease patients from the Prospective Registry in IBD study at Massachusetts General Hospital (PRISM) genotyped by CD-risk SP140 SNPs rs28445040 and rs6716753. Patient metadata is provided in **Figure S2**. Briefly, mononuclear cells were isolated by density gradient centrifugation of PBS-diluted buffy coat/blood (1:2) over Ficoll-Paque Plus (GE Healthcare). The PBMC layer was carefully removed and washed 3 times with PBS.

### Western blot

For Immunoblotting of PBMCs or human macrophages, 2.5 million cells were used and cell lysates were prepared by incubation and sonication of cells in RIPA buffer (10 mM Tris-Cl (pH 8.0), 1 mM EDTA, 0.5 mM EGTA, 1% Triton X-100, 0.1% sodium deoxycholate, 0.1% SDS, 140 mM NaCl). 20 μg protein was electrophoresed per lane on NuPAGE Novex Bis-Tris 4–12% Gels (Invitrogen) and transferred to PVDF membranes using iBlot dry blotting system (Invitrogen). The following antibodies were used: anti-S100A8 (Abcam; ab92331), anti-H3 (Abcam; ab18521).

### Quantitative PCR

RNA was extracted from cells using the RNeasy Mini Kit (Qiagen) with on-column DNase digest (Qiagen) according to manufacturer’s instructions. cDNA was synthesized from isolated RNA by reverse transcription using the iScript cDNA Synthesis Kit (Bio-Rad) according to the manufacturer’s instructions. In the case of quantitative PCR performed on stool DNA, QIAamp DNA Stool Mini Kit was used to extract DNA from feces. Quantitative PCR was performed using iTaq Universal SYBR Green Supermix (Bio-Rad) according to the manufacturer’s instructions. To analyze relative mRNA levels, derived values were normalized to indicated housekeeping genes. qPCR primers were purchased from Sigma-Aldrich. A complete list of primer sequences is provided in Supplementary Table 1.

### Bone marrow-derived macrophages

Bone marrow-derived macrophages (BMDMs) were produced from WT or Sp140-deficient mice. Bone marrow was flushed from tibia and femur and allowed to adhere to a non-treated tissue culture plate for 1 day. Non-adherent cells were then differentiated to macrophages in DMEM containing 10% FBS, 1% L-Glutamine, 1% penicillin/streptomycin, 0.1% β-mercaptoethanol, 5ng/mL of interleukin 3 (IL-3, Peprotech) and macrophage colony stimulating factor (M-CSF, Peprotech) for 7 days. Macrophage maturity was assessed by surface expression of CD11b and F4/80 with flow cytometry.

### Flow cytometry

For flow cytometric analysis, colons were dissected, and caeca were removed. The tissue was placed in RPMI containing 5% FBS and fat was removed by careful dissection followed by gentle rolling along a moist paper towel. The tissue was cut open longitudinally and rinsed to remove loose fat and fecal matter. Colon was cut into 1-2 cm pieces and transferred to RPMI containing 5% FBS, 5mM EDTA, and 1mM DTT and incubated at 37C for 20 minutes with shaking (400rpm). To remove epithelial cells, colons were next vigorously shaken in complete RPMI with 2mM EDTA. Colon was minced into fine pieces and digested in complete RPMI containing 0.5 mg/mL Dispase II (Gibco) and 1.5 mg/mL type IV collagenase (Gibco) for 45 minutes at 37C shaking (400rpm). Single-cell suspensions were prepared using a 40um nylon cell strainer. Cells were stained with fixable live/dead stain, Fc Block, counting beads, and fluorochrome-conjugated antibodies for 20 min on ice then fixed on ice for 30 min using Fixation/Permeabilization Solution Kit. Data were acquired using a BD FACSAria running FACSDiva software and analyzed with FlowJo software. Mononuclear phagocytes were analyzed using a previously described gating strategy (Adiliaghdam et al., 2021). See Supplemental Table 2 for a list of FACS antibodies and reagents used.

### Mouse bone marrow reconstitution

Bone marrow progenitor cells obtained from C57BL/6 donor mice were placed in 24-well tissue culture dishes (1 × 10^6^ cells per well) in fresh media containing 15% fetal bovine serum, IL-3 (20 ng/ml), IL-6 (50 ng/ml), and stem cell factor (SCF) (50 ng/ml). On day 4, the cells were harvested and washed twice in PBS. For bone marrow transplantation, C57BL/6 recipient mice were irradiated with 131Cs at 10 gray on day 0 and injected retro-orbitally with 2.5 × 10^6^ bone marrow cells in 200 μl of Dulbecco’s modified Eagle’s medium, after which the mice were monitored for 6 weeks to allow for immune reconstitution.

### Dextran sodium sulfate (DSS)-colitis

Mice were administered 2% dextran sodium sulfate salt (DSS, MW = 36,000-50,000 Da; MP Biomedicals) in drinking water ad libitum for 7 days (freshly prepared every other day), followed by regular drinking water for 5 days. Mice were sacrificed on day 12. Colon length was measured. For histology, tissue was collected from the distal part of the colon, and hematoxylin and eosin staining were performed. Images were obtained on an EVOS XL Core Imaging System microscope (ThermoFisher). Metronizadole treated mice were administered 1g/L metronidazole drinking water with 8mg/mL Sweet’N Low for 2 weeks prior to the start of DSS administration and continued through day 12.

### 16S rRNA gene sequencing

QIAamp DNA Stool Mini Kit was used to extract DNA from mouse and human feces. The hypervariable region (V4) of the 16S rRNA gene sequences were amplified by using PCR with adaptor primer set containing 12bp multiplex identifier sequences (Integrated DNA Technologies, Coralville, IA). Each reaction contained template DNA, each primer (0.2 μM), and 2× Platinum™ Hot Start PCR Master Mix (Invitrogen, Carlsbad, CA). PCR conditions were as follows: 94°C for 3 min, followed by 22 cycles at 94°C for 45 sec, 52°C for 1 min, and 72°C for 1.5 min and a final extension step at 72°C for 10 min. Triplicate reactions were prepared, pooled, and purified using SpeedBead™ carboxylate magnetic bead solution (GE Healthcare, Marlborough, MA). The amplicons were quantified using KAPA Library Quantitation kit (KAPA, Cape Town, South Africa) and an equimolar amount of each sample was sequenced on the MiSeq System (Illumina, San Diego, CA) with 2×150 paired-end run parameters. Microbiome bioinformatics analysis was performed with QIIME 2 release 2018.11. In brief, cutadapt was used to trim 515F/806R primers in the paired end sequences. Subsequently, the sequences were stripped of noise by using the DADA2 denoiser program. The denoised, trimmed and higher quality amplicon sequence variants (ASVs) were aligned with MAFFT and used to construct a phylogeny with fasttree2. All ASVs were assigned taxonomy using the q2-feature-classifier. In brief, a naïve Bayes classifier was trained on the Greengenes 13_8 99% OTUs reference sequences to assign taxonomy to each ASV. The compositional differences among the groups were determined by a linear discriminant analysis using LEfSe (Segata et al., 2011) with a threshold of 2.0 on the logarithmic score and alpha value of 0.01 for the factorial Kruskal-Wallis test among classes using Galaxy (https://huttenhower.sph.harvard.edu/galaxy/). 16S rRNA sequencing data are deposited at Sequencing Read Archive (SRA) under accession number SAMN26873569.

### Macrophage stimulations with stool homogenates

Stool pellets were weighed and resuspended in 40mg/mL complete DMEM then homogenized using a BeadBug 3 microtube homogenizer (Benchmark Scientific) for 9 minutes at highest speed in BeadBug tubes containing 1.5mm Zirconium beads. Homogenized stool was centrifuged at 400g for 1 min. Supernatant was collected and passed through 70μm strainer. Stool homogenates were then added to 0.5×10^6^ bone marrow-derived macrophages (BMDMs) in a 96 well plate at a final concentration of 1mg/mL. Antibiotic treated mice were administered a broad-spectrum antibiotic cocktail (1g/L metronidazole, 0.5g/L vancomycin, 1g/L ampicillin, 1g/L neomycin) containing 8mg/mL Sweet’N Low via drinking water. Stool was collected after 4 weeks of antibiotic treatment. Mouse IL-6 was measured in cell supernatants using a DuoSet ELISA kit per manufacturer instructions (R&D Systems).

### Software

Flow cytometry samples were acquired using FACSDiva and analyzed using FlowJo version 10. Statistical analysis was performed using GraphPad Prism version 7.

## References

Adiliaghdam, F., H. Amatullah, S. Digumarthi, T.L. Saunders, R.-U. Rahman, L.P. Wong, R. Sadreyev, L. Droit, J. Paquette, P. Goyette, J. Rioux, R. Hodin, K.A. Mihindukulasuriya, S.A. Handley, and K.L. Jeffrey. 2021. Human enteric viruses shape disease phenotype through divergent innate immunomodulation. bioRxiv. doi:10.1101/2021.10.14.464404.

Alam, A., G. Leoni, M. Quiros, H. Wu, C. Desai, H. Nishio, R.M. Jones, A. Nusrat, and A.S. Neish. 2016. The microenvironment of injured murine gut elicits a local pro-restitutive microbiota. Nature Microbiology 1:15021.

Amatullah, H., S. Digumarthi, I. Fraschilla, F. Adiliaghdam, G. Bonilla, L.P. Wong, R.I. Sadreyev, and K.L. Jeffrey. 2021. Identification of topoisomerase as a precision-medicine target in chromatin reader SP140-driven Crohn’s disease. 10.1101/2021.09.20.461083.

Amatullah, H., and K.L. Jeffrey. 2020. Epigenome-metabolome-microbiome axis in health and IBD. Curr Opin Microbiol 56:97–108.

Ansaldo, E., L.C. Slayden, K.L. Ching, M.A. Koch, N.K. Wolf, D.R. Plichta, E.M. Brown, D.B. Graham, R.J. Xavier, J.J. Moon, and G.M. Barton. 2019. Akkermansia muciniphila induces intestinal adaptive immune responses during homeostasis. Science 364:1179–1184.

Argiropoulos, B., and R.K. Humphries. 2007. Hox genes in hematopoiesis and leukemogenesis. Oncogene 26:6766–6776.

Arrowsmith, C.H., C. Bountra, P.V. Fish, K. Lee, and M. Schapira. 2012. Epigenetic protein families: a new frontier for drug discovery. Nat Rev Drug Discov 11:384–400.

Bain, C.C., A. Bravo-Blas, C.L. Scott, E. Gomez Perdiguero, F. Geissmann, S. Henri, B. Malissen, L.C. Osborne, D. Artis, and A.M. Mowat. 2014. Constant replenishment from circulating monocytes maintains the macrophage pool in the intestine of adult mice. Nature Immunology 15:929–937.

Berer, K., L.A. Gerdes, E. Cekanaviciute, X. Jia, L. Xiao, Z. Xia, C. Liu, L. Klotz, U. Stauffer, S.E. Baranzini, T. Kümpfel, R. Hohlfeld, G. Krishnamoorthy, and H. Wekerle. 2017. Gut microbiota from multiple sclerosis patients enables spontaneous autoimmune encephalomyelitis in mice. Proceedings of the National Academy of Sciences 114:10719–10724.

Berer, K., M. Mues, M. Koutrolos, Z.A. Rasbi, M. Boziki, C. Johner, H. Wekerle, and G. Krishnamoorthy. 2011. Commensal microbiota and myelin autoantigen cooperate to trigger autoimmune demyelination. Nature 479:538–541.

Bienz, M. 2006. The PHD finger, a nuclear protein-interaction domain. Trends in Biochemical Sciences 31:35–40.

Blander, J.M., R.S. Longman, I.D. Iliev, G.F. Sonnenberg, and D. Artis. 2017. Regulation of inflammation by microbiota interactions with the host. Nature Immunology 18:851–860.

Bottomley, M.J., M.W. Collard, J.I. Huggenvik, Z. Liu, T.J. Gibson, and M. Sattler. 2001. Nature Structural Biology 8:626–633.

Britton, G.J., E.J. Contijoch, I. Mogno, O.H. Vennaro, S.R. Llewellyn, R. Ng, Z. Li, A. Mortha, M. Merad, A. Das, D. Gevers, D.P.B. McGovern, N. Singh, J. Braun, J.P. Jacobs, J.C. Clemente, A. Grinspan, B.E. Sands, J.-F. Colombel, M.C. Dubinsky, and J.J. Faith. 2019. Microbiotas from Humans with Inflammatory Bowel Disease Alter the Balance of Gut Th17 and RORγt+ Regulatory T Cells and Exacerbate Colitis in Mice. Immunity 50:212–224.e214.

Britton, G.J., E.J. Contijoch, M.P. Spindler, V. Aggarwala, B. Dogan, G. Bongers, L. San Mateo, A. Baltus, A. Das, D. Gevers, T.J. Borody, N.O. Kaakoush, M.A. Kamm, H. Mitchell, S. Paramsothy, J.C. Clemente, J.F. Colombel, K.W. Simpson, M.C. Dubinsky, A. Grinspan, and J.J. Faith. 2020. Defined microbiota transplant restores Th17/RORgammat(+) regulatory T cell balance in mice colonized with inflammatory bowel disease microbiotas. Proc Natl Acad Sci U S A 117:21536–21545.

Cahill, R.J., C.J. Foltz, J.G. Fox, C.A. Dangler, F. Powrie, and D.B. Schauer. 1997. Inflammatory bowel disease: an immunity-mediated condition triggered by bacterial infection with Helicobacter hepaticus. Infect Immun 65:3126–3131.

Caruso, R., T. Mathes, E.C. Martens, N. Kamada, A. Nusrat, N. Inohara, and G. Nunez. 2019. A specific gene-microbe interaction drives the development of Crohn’s disease-like colitis in mice. Sci Immunol. doi:10.1126/sciimmunol.aaw4341.

Cekanaviciute, E., B.B. Yoo, T.F. Runia, J.W. Debelius, S. Singh, C.A. Nelson, R. Kanner, Y. Bencosme, Y.K. Lee, S.L. Hauser, E. Crabtree-Hartman, I.K. Sand, M. Gacias, Y. Zhu, P. Casaccia, B.A.C. Cree, R. Knight, S.K. Mazmanian, and S.E. Baranzini. 2017. Gut bacteria from multiple sclerosis patients modulate human T cells and exacerbate symptoms in mouse models. Proceedings of the National Academy of Sciences 114:10713–10718.

Chai, J.N., Y. Peng, S. Rengarajan, B.D. Solomon, T.L. Ai, Z. Shen, J.S.A. Perry, K.A. Knoop, T. Tanoue, S. Narushima, K. Honda, C.O. Elson, R.D. Newberry, T.S. Stappenbeck, A.L. Kau, D.A. Peterson, J.G. Fox, and C.S. Hsieh. 2017. Helicobacter species are potent drivers of colonic T cell responses in homeostasis and inflammation. Sci Immunol. doi:10.1126/sciimmunol.aal5068.

Conway, K.L., P. Kuballa, J.H. Song, K.K. Patel, A.B. Castoreno, O.H. Yilmaz, H.B. Jijon, M. Zhang, L.N. Aldrich, E.J. Villablanca, J.M. Peloquin, G. Goel, I.A. Lee, E. Mizoguchi, H.N. Shi, A.K. Bhan, S.Y. Shaw, S.L. Schreiber, H.W. Virgin, A.F. Shamji, T.S. Stappenbeck, H.C. Reinecker, and R.J. Xavier. 2013. Atg16l1 is Required for Autophagy in Intestinal Epithelial Cells and Protection of Mice From Salmonella Infection. Gastroenterology 145:1347–1357.

Corpet, A., C. Kleijwegt, S. Roubille, F. Juillard, K. Jacquet, P. Texier, and P. Lomonte. 2020. PML nuclear bodies and chromatin dynamics: catch me if you can! Nucleic Acids Research 48:11890–11912.

De Santa, F., M.G. Totaro, E. Prosperini, S. Notarbartolo, G. Testa, and G. Natoli. 2007. The Histone H3 Lysine-27 Demethylase Jmjd3 Links Inflammation to Inhibition of Polycomb-Mediated Gene Silencing. Cell 130:1083–1094.

Denson, L.A., I. Jurickova, R. Karns, K.A. Shaw, D.J. Cutler, D.T. Okou, A. Dodd, K. Quinn, K. Mondal, B.J. Aronow, Y. Haberman, A. Linn, A. Price, R. Bezold, K. Lake, K. Jackson, T.D. Walters, A. Griffiths, R.N. Baldassano, J.D. Noe, J.S. Hyams, W.V. Crandall, B.S. Kirschner, M.B. Heyman, S. Snapper, S.L. Guthery, M.C. Dubinsky, N.S. Leleiko, A.R. Otley, R.J. Xavier, C. Stevens, M.J. Daly, M.E. Zwick, and S. Kugathasan. 2018. Clinical and Genomic Correlates of Neutrophil Reactive Oxygen Species Production in Pediatric Patients With Crohn’s Disease. Gastroenterology 154:2097–2110.

Elinav, E., T. Strowig, Andrew, J. Henao-Mejia, Christoph, Carmen, David, J. Bertin, Stephanie, Jeffrey, and Richard. 2011. NLRP6 Inflammasome Regulates Colonic Microbial Ecology and Risk for Colitis. Cell 145:745–757.

Feng, T., L. Wang, T.R. Schoeb, C.O. Elson, and Y. Cong. 2010. Microbiota innate stimulation is a prerequisite for T cell spontaneous proliferation and induction of experimental colitis. Journal of Experimental Medicine 207:1321–1332.

Filippakopoulos, P., S. Picaud, M. Mangos, T. Keates, J.P. Lambert, D. Barsyte-Lovejoy, I. Felletar, R. Volkmer, S. Muller, T. Pawson, A.C. Gingras, C.H. Arrowsmith, and S. Knapp. 2012. Histone recognition and large-scale structural analysis of the human bromodomain family. Cell 149:214–231.

Franzosa, E.A., A. Sirota-Madi, J. Avila-Pacheco, N. Fornelos, H.J. Haiser, S. Reinker, T. Vatanen, A.B. Hall, H. Mallick, L.J. McIver, J.S. Sauk, R.G. Wilson, B.W. Stevens, J.M. Scott, K. Pierce, A.A. Deik, K. Bullock, F. Imhann, J.A. Porter, A. Zhernakova, J. Fu, R.K. Weersma, C. Wijmenga, C.B. Clish, H. Vlamakis, C. Huttenhower, and R.J. Xavier. 2019. Gut microbiome structure and metabolic activity in inflammatory bowel disease. Nature Microbiology 4:293–305.

Fraschilla, I., and K.L. Jeffrey. 2020. The Speckled Protein (SP) Family: Immunity’s Chromatin Readers. Trends in Immunology 41:572–585.

Frederic, O. Koren, Julia, Malin, I. Nalbantoglu, Jesse, Y. Su, B. Chassaing, William, A. González, Jose, Tyler, N. Barnich, A. Darfeuille-Michaud, M. Vijay-Kumar, R. Knight, Ruth, and Andrew. 2012. Transient Inability to Manage Proteobacteria Promotes Chronic Gut Inflammation in TLR5-Deficient Mice. Cell Host & Microbe 12:139–152.

Garcia-Dominguez, M., R. March-Diaz, and J.C. Reyes. 2008. The PHD Domain of Plant PIAS Proteins Mediates Sumoylation of Bromodomain GTE Proteins. Journal of Biological Chemistry 283:21469–21477.

Garrett, W.S., C.A. Gallini, T. Yatsunenko, M. Michaud, A. Dubois, M.L. Delaney, S. Punit, M. Karlsson, L. Bry, J.N. Glickman, J.I. Gordon, A.B. Onderdonk, and L.H. Glimcher. 2010. Enterobacteriaceae Act in Concert with the Gut Microbiota to Induce Spontaneous and Maternally Transmitted Colitis. Cell Host & Microbe 8:292–300.

Graham, D.B., and R.J. Xavier. 2020. Pathway paradigms revealed from the genetics of inflammatory bowel disease. Nature 578:527–539.

Hoischen, C., S. Monajembashi, K. Weisshart, and P. Hemmerich. 2018. Multimodal Light Microscopy Approaches to Reveal Structural and Functional Properties of Promyelocytic Leukemia Nuclear Bodies. Front Oncol 8:125.

Huang, Y., K. Sitwala, J. Bronstein, D. Sanders, M. Dandekar, C. Collins, G. Robertson, J. Macdonald, T. Cezard, M. Bilenky, N. Thiessen, Y. Zhao, T. Zeng, M. Hirst, A. Hero, S. Jones, and J.L. Hess. 2012. Identification and characterization of Hoxa9 binding sites in hematopoietic cells. Blood 119:388–398.

Huoh, Y.-S., B. Wu, S. Park, D. Yang, K. Bansal, E. Greenwald, W.P. Wong, D. Mathis, and S. Hur. 2020. Dual functions of Aire CARD multimerization in the transcriptional regulation of T cell tolerance. Nature Communications. doi:10.1038/s41467-020-15448-w.

Iliev, I.D., and K. Cadwell. 2021. Effects of Intestinal Fungi and Viruses on Immune Responses and Inflammatory Bowel Diseases. Gastroenterology 160:1050–1066.

International Multiple Sclerosis Genetics, C., A.H. Beecham, N.A. Patsopoulos, D.K. Xifara, M.F. Davis, A. Kemppinen, C. Cotsapas, T.S. Shah, C. Spencer, D. Booth, A. Goris, A. Oturai, J. Saarela, B. Fontaine, B. Hemmer, C. Martin, F. Zipp, S. D’Alfonso, F. Martinelli-Boneschi, B. Taylor, H.F. Harbo, I. Kockum, J. Hillert, T. Olsson, M. Ban, J.R. Oksenberg, R. Hintzen, L.F. Barcellos, C. Wellcome Trust Case Control, I.B.D.G.C. International, C. Agliardi, L. Alfredsson, M. Alizadeh, C. Anderson, R. Andrews, H.B. Sondergaard, A. Baker, G. Band, S.E. Baranzini, N. Barizzone, J. Barrett, C. Bellenguez, L. Bergamaschi, L. Bernardinelli, A. Berthele, V. Biberacher, T.M. Binder, H. Blackburn, I.L. Bomfim, P. Brambilla, S. Broadley, B. Brochet, L. Brundin, D. Buck, H. Butzkueven, S.J. Caillier, W. Camu, W. Carpentier, P. Cavalla, E.G. Celius, I. Coman, G. Comi, L. Corrado, L. Cosemans, I. Cournu-Rebeix, B.A. Cree, D. Cusi, V. Damotte, G. Defer, S.R. Delgado, P. Deloukas, A. di Sapio, A.T. Dilthey, P. Donnelly, B. Dubois, M. Duddy, S. Edkins, I. Elovaara, F. Esposito, N. Evangelou, B. Fiddes, J. Field, A. Franke, C. Freeman, I.Y. Frohlich, D. Galimberti, C. Gieger, P.A. Gourraud, C. Graetz, A. Graham, V. Grummel, C. Guaschino, A. Hadjixenofontos, H. Hakonarson, C. Halfpenny, G. Hall, P. Hall, A. Hamsten, J. Harley, T. Harrower, C. Hawkins, G. Hellenthal, C. Hillier, J. Hobart, M. Hoshi, S.E. Hunt, M. Jagodic, I. Jelcic, A. Jochim, B. Kendall, A. Kermode, T. Kilpatrick, K. Koivisto, I. Konidari, T. Korn, H. Kronsbein, C. Langford, M. Larsson, M. Lathrop, C. Lebrun-Frenay, J. Lechner-Scott, M.H. Lee, M.A. Leone, V. Leppa, G. Liberatore, B.A. Lie, C.M. Lill, M. Linden, J. Link, F. Luessi, J. Lycke, F. Macciardi, S. Mannisto, C.P. Manrique, R. Martin, V. Martinelli, D. Mason, G. Mazibrada, C. McCabe, I.L. Mero, J. Mescheriakova, L. Moutsianas, K.M. Myhr, G. Nagels, R. Nicholas, P. Nilsson, F. Piehl, M. Pirinen, S.E. Price, H. Quach, M. Reunanen, W. Robberecht, N.P. Robertson, M. Rodegher, D. Rog, M. Salvetti, N.C. Schnetz-Boutaud, F. Sellebjerg, R.C. Selter, C. Schaefer, S. Shaunak, L. Shen, S. Shields, V. Siffrin, M. Slee, P.S. Sorensen, M. Sorosina, M. Sospedra, A. Spurkland, A. Strange, E. Sundqvist, V. Thijs, J. Thorpe, A. Ticca, P. Tienari, C. van Duijn, E.M. Visser, S. Vucic, H. Westerlind, J.S. Wiley, A. Wilkins, J.F. Wilson, J. Winkelmann, J. Zajicek, E. Zindler, J.L. Haines, M.A. Pericak-Vance, A.J. Ivinson, G. Stewart, D. Hafler, S.L. Hauser, A. Compston, G. McVean, P. De Jager, S.J. Sawcer, and J.L. McCauley. 2013. Analysis of immune-related loci identifies 48 new susceptibility variants for multiple sclerosis. Nat Genet 45:1353–1360.

Ivanov, A.V., H. Peng, V. Yurchenko, K.L. Yap, D.G. Negorev, D.C. Schultz, E. Psulkowski, W.J. Fredericks, D.E. White, G.G. Maul, M.J. Sadofsky, M.M. Zhou, and F.J. Rauscher, 3rd. 2007. PHD domain-mediated E3 ligase activity directs intramolecular sumoylation of an adjacent bromodomain required for gene silencing. Mol Cell 28:823–837.

Ji, D.X., K.C. Witt, D.I. Kotov, S.R. Margolis, A. Louie, V. Chevée, K.J. Chen, M. Gaidt, H.S. Dhaliwal, A.Y. Lee, S.L. Nishimura, D.S. Zamboni, I. Kramnik, D.A. Portnoy, K.H. Darwin, and R.E. Vance. 2021. Role of the transcriptional regulator SP140 in resistance to bacterial infections via repression of type I interferons. eLife. doi:10.7554/elife.67290.

Jostins, L., S. Ripke, R.K. Weersma, R.H. Duerr, D.P. McGovern, K.Y. Hui, J.C. Lee, L.P. Schumm, Y. Sharma, C.A. Anderson, J. Essers, M. Mitrovic, K. Ning, I. Cleynen, E. Theatre, S.L. Spain, S. Raychaudhuri, P. Goyette, Z. Wei, C. Abraham, J.P. Achkar, T. Ahmad, L. Amininejad, A.N. Ananthakrishnan, V. Andersen, J.M. Andrews, L. Baidoo, T. Balschun, P.A. Bampton, A. Bitton, G. Boucher, S. Brand, C. Buning, A. Cohain, S. Cichon, M. D’Amato, D. De Jong, K.L. Devaney, M. Dubinsky, C. Edwards, D. Ellinghaus, L.R. Ferguson, D. Franchimont, K. Fransen, R. Gearry, M. Georges, C. Gieger, J. Glas, T. Haritunians, A. Hart, C. Hawkey, M. Hedl, X. Hu, T.H. Karlsen, L. Kupcinskas, S. Kugathasan, A. Latiano, D. Laukens, I.C. Lawrance, C.W. Lees, E. Louis, G. Mahy, J. Mansfield, A.R. Morgan, C. Mowat, W. Newman, O. Palmieri, C.Y. Ponsioen, U. Potocnik, N.J. Prescott, M. Regueiro, J.I. Rotter, R.K. Russell, J.D. Sanderson, M. Sans, J. Satsangi, S. Schreiber, L.A. Simms, J. Sventoraityte, S.R. Targan, K.D. Taylor, M. Tremelling, H.W. Verspaget, M. De Vos, C. Wijmenga, D.C. Wilson, J. Winkelmann, R.J. Xavier, S. Zeissig, B. Zhang, C.K. Zhang, H. Zhao, I.B.D.G.C. International, M.S. Silverberg, V. Annese, H. Hakonarson, S.R. Brant, G. Radford-Smith, C.G. Mathew, J.D. Rioux, E.E. Schadt, M.J. Daly, A. Franke, M. Parkes, S. Vermeire, J.C. Barrett, and J.H. Cho. 2012. Host-microbe interactions have shaped the genetic architecture of inflammatory bowel disease. Nature 491:119–124.

Kaiko, G.E., S.H. Ryu, O.I. Koues, P.L. Collins, L. Solnica-Krezel, E.J. Pearce, E.L. Pearce, E.M. Oltz, and T.S. Stappenbeck. 2016. The Colonic Crypt Protects Stem Cells from Microbiota-Derived Metabolites. Cell 165:1708–1720.

Knights, D., M.S. Silverberg, R.K. Weersma, D. Gevers, G. Dijkstra, H. Huang, A.D. Tyler, S. van Sommeren, F. Imhann, J.M. Stempak, H. Huang, P. Vangay, G.A. Al-Ghalith, C. Russell, J. Sauk, J. Knight, M.J. Daly, C. Huttenhower, and R.J. Xavier. 2014. Complex host genetics influence the microbiome in inflammatory bowel disease. Genome Med 6:107.

Kugelberg, E. 2014. Curbing gut inflammation. Nature Reviews Immunology 14:583–583.

Kullberg, M.C., J.M. Ward, P.L. Gorelick, P. Caspar, S. Hieny, A. Cheever, D. Jankovic, and A. Sher. 1998. Helicobacter hepaticus triggers colitis in specific-pathogen-free interleukin-10 (IL-10)-deficient mice through an IL-12-and gamma interferon-dependent mechanism. Infect Immun 66:5157–5166.

Liu, J.Z., S. Van Sommeren, H. Huang, S.C. Ng, R. Alberts, A. Takahashi, S. Ripke, J.C. Lee, L. Jostins, T. Shah, S. Abedian, J.H. Cheon, J. Cho, N.E. Daryani, L. Franke, Y. Fuyuno, A. Hart, R.C. Juyal, G. Juyal, W.H. Kim, A.P. Morris, H. Poustchi, W.G. Newman, V. Midha, T.R. Orchard, H. Vahedi, A. Sood, J.J.Y. Sung, R. Malekzadeh, H.-J. Westra, K. Yamazaki, S.-K. Yang, J.C. Barrett, A. Franke, B.Z. Alizadeh, M. Parkes, T. B K, M.J. Daly, M. Kubo, C.A. Anderson, and R.K. Weersma. 2015. Association analyses identify 38 susceptibility loci for inflammatory bowel disease and highlight shared genetic risk across populations. Nature Genetics 47:979–986.

Lloyd-Price, J., C. Arze, A.N. Ananthakrishnan, M. Schirmer, J. Avila-Pacheco, T.W. Poon, E. Andrews, N.J. Ajami, K.S. Bonham, C.J. Brislawn, D. Casero, H. Courtney, A. Gonzalez, T.G. Graeber, A.B. Hall, K. Lake, C.J. Landers, H. Mallick, D.R. Plichta, M. Prasad, G. Rahnavard, J. Sauk, D. Shungin, Y. Vázquez-Baeza, R.A. White, J. Braun, L.A. Denson, J.K. Jansson, R. Knight, S. Kugathasan, D.P.B. McGovern, J.F. Petrosino, T.S. Stappenbeck, H.S. Winter, C.B. Clish, E.A. Franzosa, H. Vlamakis, R.J. Xavier, and C. Huttenhower. 2019. Multi-omics of the gut microbial ecosystem in inflammatory bowel diseases. Nature 569:655–662.

Madani, N., R. Millette, E.J. Platt, M. Marin, S.L. Kozak, D.B. Bloch, and D. Kabat. 2002. Implication of the Lymphocyte-Specific Nuclear Body Protein Sp140 in an Innate Response to Human Immunodeficiency Virus Type 1. Journal of Virology 76:11133–11138.

Matesanz, F., V. Potenciano, M. Fedetz, P. Ramos-Mozo, M. Abad-Grau Mdel, M. Karaky, C. Barrionuevo, G. Izquierdo, J.L. Ruiz-Pena, M.I. Garcia-Sanchez, M. Lucas, O. Fernandez, L. Leyva, D. Otaegui, M. Munoz-Culla, J. Olascoaga, K. Vandenbroeck, I. Alloza, I. Astobiza, A. Antiguedad, L.M. Villar, J.C. Alvarez-Cermeno, S. Malhotra, M. Comabella, X. Montalban, A. Saiz, Y. Blanco, R. Arroyo, J. Varade, E. Urcelay, and A. Alcina. 2015. A functional variant that affects exon-skipping and protein expression of SP140 as genetic mechanism predisposing to multiple sclerosis. Hum Mol Genet 24:5619–5627.

Matsushita, K., X. Li, Y. Nakamura, D. Dong, K. Mukai, M. Tsai, S.B. Montgomery, and S.J. Galli. 2021. The role of Sp140 revealed in IgE and mast cell responses in Collaborative Cross mice. JCI Insight 6:2379–3708.

Mehta, S., D.A. Cronkite, M. Basavappa, T.L. Saunders, F. Adiliaghdam, H. Amatullah, S.A. Morrison, J.D. Pagan, R.M. Anthony, P. Tonnerre, G.M. Lauer, J.C. Lee, S. Digumarthi, L. Pantano, S.J. Ho Sui, F. Ji, R. Sadreyev, C. Zhou, A.C. Mullen, V. Kumar, Y. Li, C. Wijmenga, R.J. Xavier, T.K. Means, and K.L. Jeffrey. 2017. Maintenance of macrophage transcriptional programs and intestinal homeostasis by epigenetic reader SP140. Sci Immunol. doi:10.1126/sciimmunol.aag3160.

Morgan, X.C., T.L. Tickle, H. Sokol, D. Gevers, K.L. Devaney, D.V. Ward, J.A. Reyes, S.A. Shah, N. LeLeiko, S.B. Snapper, A. Bousvaros, J. Korzenik, B.E. Sands, R.J. Xavier, and C. Huttenhower. 2012. Dysfunction of the intestinal microbiome in inflammatory bowel disease and treatment. Genome Biol 13:R79.

Muise, A.M., W. Xu, C.-H. Guo, T.D. Walters, V.M. Wolters, R. Fattouh, G.Y. Lam, P. Hu, R. Murchie, M. Sherlock, J.C. Gana, R.K. Russell, M. Glogauer, R.H. Duerr, J.H. Cho, C.W. Lees, J. Satsangi, D.C. Wilson, A.D. Paterson, A.M. Griffiths, M.S. Silverberg, and J.H. Brumell. 2012. NADPH oxidase complex and IBD candidate gene studies: identification of a rare variant in NCF2 that results in reduced binding to RAC2. Gut 61:1028–1035.

Ni, J., T.D. Shen, E.Z. Chen, K. Bittinger, A. Bailey, M. Roggiani, A. Sirota-Madi, E.S. Friedman, L. Chau, A. Lin, I. Nissim, J. Scott, A. Lauder, C. Hoffmann, G. Rivas, L. Albenberg, R.N. Baldassano, J. Braun, R.J. Xavier, C.B. Clish, M. Yudkoff, H. Li, M. Goulian, F.D. Bushman, J.D. Lewis, and G.D. Wu. 2017. A role for bacterial urease in gut dysbiosis and Crohn’s disease. Sci Transl Med. doi:10.1126/scitranslmed.aah6888.

Olbjørn, C., M. Cvancarova Småstuen, E. Thiis-Evensen, B. Nakstad, M.H. Vatn, J. Jahnsen, P. Ricanek, S. Vatn, A.E. Moen, T.M. Tannæs, J.C. Lindstrøm, J.D. Söderholm, J. Halfvarson, F. Gomollón, C. Casén, M.K. Karlsson, R. Kalla, A.T. Adams, J. Satsangi, and G. Perminow. 2019. Fecal microbiota profiles in treatment-naive pediatric inflammatory bowel disease - associations with disease phenotype, treatment, and outcome. Clinical and Experimental Gastroenterology Volume 12:37–49.

Pan, H., B.-S. Yan, M. Rojas, Y.V. Shebzukhov, H. Zhou, L. Kobzik, D.E. Higgins, M.J. Daly, B.R. Bloom, and I. Kramnik. 2005. Ipr1 gene mediates innate immunity to tuberculosis. Nature 434:767–772.

Peng, J., and J. Wysocka. 2008. It takes a PHD to SUMO. Trends Biochem Sci 33:191–194.

Pichugin, A.V., B.S. Yan, A. Sloutsky, L. Kobzik, and I. Kramnik. 2009. Dominant role of the sst1 locus in pathogenesis of necrotizing lung granulomas during chronic tuberculosis infection and reactivation in genetically resistant hosts. Am J Pathol 174:2190–2201.

Plichta, D.R., D.B. Graham, S. Subramanian, and R.J. Xavier. 2019. Therapeutic Opportunities in Inflammatory Bowel Disease: Mechanistic Dissection of Host-Microbiome Relationships. Cell 178:1041–1056.

Ramanan, D., M.S. Tang, R. Bowcutt, P. Loke, and K. Cadwell. 2014. Bacterial sensor Nod2 prevents inflammation of the small intestine by restricting the expansion of the commensal Bacteroides vulgatus. Immunity 41:311–324.

Regad, T., and M.K. Chelbi-Alix. 2001. Role and fate of PML nuclear bodies in response to interferon and viral infections. Oncogene 20:7274–7286.

Rivas, M.A., M. Beaudoin, A. Gardet, C. Stevens, Y. Sharma, C.K. Zhang, G. Boucher, S. Ripke, D. Ellinghaus, N. Burtt, T. Fennell, A. Kirby, A. Latiano, P. Goyette, T. Green, J. Halfvarson, T. Haritunians, J.M. Korn, F. Kuruvilla, C. Lagacé, B. Neale, K.S. Lo, P. Schumm, L. Törkvist, M.C. Dubinsky, S.R. Brant, M.S. Silverberg, R.H. Duerr, D. Altshuler, S. Gabriel, G. Lettre, A. Franke, M. D’Amato, D.P.B. McGovern, J.H. Cho, J.D. Rioux, R.J. Xavier, and M.J. Daly. 2011. Deep resequencing of GWAS loci identifies independent rare variants associated with inflammatory bowel disease. Nature Genetics 43:1066–1073.

Schulthess, J., S. Pandey, M. Capitani, K.C. Rue-Albrecht, I. Arnold, F. Franchini, A. Chomka, N.E. Ilott, D.G.W. Johnston, E. Pires, J. McCullagh, S.N. Sansom, C.V. Arancibia-Cárcamo, H.H. Uhlig, and F. Powrie. 2019. The Short Chain Fatty Acid Butyrate Imprints an Antimicrobial Program in Macrophages. Immunity 50:432–445.e437.

Segata, N., J. Izard, L. Waldron, D. Gevers, L. Miropolsky, W.S. Garrett, and C. Huttenhower. 2011. Metagenomic biomarker discovery and explanation. Genome Biology 12:R60.

Smythies, L.E., M. Sellers, R.H. Clements, M. Mosteller-Barnum, G. Meng, W.H. Benjamin, J.M. Orenstein, and P.D. Smith. 2005. Human intestinal macrophages display profound inflammatory anergy despite avid phagocytic and bacteriocidal activity. 115:66–75.

Stappenbeck, T.S., and H.W. Virgin. 2016. Accounting for reciprocal host-microbiome interactions in experimental science. Nature 534:191–199.

Tanoue, T., S. Morita, D.R. Plichta, A.N. Skelly, W. Suda, Y. Sugiura, S. Narushima, H. Vlamakis, I. Motoo, K. Sugita, A. Shiota, K. Takeshita, K. Yasuma-Mitobe, D. Riethmacher, T. Kaisho, J.M. Norman, D. Mucida, M. Suematsu, T. Yaguchi, V. Bucci, T. Inoue, Y. Kawakami, B. Olle, B. Roberts, M. Hattori, R.J. Xavier, K. Atarashi, and K. Honda. 2019. A defined commensal consortium elicits CD8 T cells and anti-cancer immunity. Nature 565:600–605.

Thorsteinsdottir, U., A. Mamo, E. Kroon, L. Jerome, J. Bijl, H.J. Lawrence, K. Humphries, and G. Sauvageau. 2002. Overexpression of the myeloid leukemia-associatedHoxa9 gene in bone marrow cells induces stem cell expansion. Blood 99:121–129.

Vinolo, M.A.R., H.G. Rodrigues, R.T. Nachbar, and R. Curi. 2011. Regulation of Inflammation by Short Chain Fatty Acids. Nutrients 3:858–876.

Walujkar, S.A., D.P. Dhotre, N.P. Marathe, P.S. Lawate, R.S. Bharadwaj, and Y.S. Shouche. 2014. Characterization of bacterial community shift in human Ulcerative Colitis patients revealed by Illumina based 16S rRNA gene amplicon sequencing. Gut Pathogens 6:22.

Waterfield, M., I.S. Khan, J.T. Cortez, U. Fan, T. Metzger, A. Greer, K. Fasano, M. Martinez-Llordella, J.L. Pollack, D.J. Erle, M. Su, and M.S. Anderson. 2014. The transcriptional regulator Aire coopts the repressive ATF7ip-MBD1 complex for the induction of immunotolerance. Nat Immunol 15:258–265.

Wu, S.E., S. Hashimoto-Hill, V. Woo, E.M. Eshleman, J. Whitt, L. Engleman, R. Karns, L.A. Denson, D.B. Haslam, and T. Alenghat. 2020. Microbiota-derived metabolite promotes HDAC3 activity in the gut. Nature 586:108–112.

Yang, I., D. Eibach, F. Kops, B. Brenneke, S. Woltemate, J. Schulze, A. Bleich, A.D. Gruber, S. Muthupalani, J.G. Fox, C. Josenhans, and S. Suerbaum. 2013. Intestinal Microbiota Composition of Interleukin-10 Deficient C57BL/6J Mice and Susceptibility to Helicobacter hepaticus-Induced Colitis. PLoS ONE 8:e70783.

Yu, A.I., L. Zhao, K.A. Eaton, S. Ho, J. Chen, S. Poe, J. Becker, A. Gonzalez, D. McKinstry, M. Hasso, J. Mendoza-Castrejon, J. Whitfield, C. Koumpouras, P.D. Schloss, E.C. Martens, and G.Y. Chen. 2020. Gut Microbiota Modulate CD8 T Cell Responses to Influence Colitis-Associated Tumorigenesis. Cell Reports 31:107471.

Zenewicz, L.A., X. Yin, G. Wang, E. Elinav, L. Hao, L. Zhao, and R.A. Flavell. 2013. IL-22 Deficiency Alters Colonic Microbiota To Be Transmissible and Colitogenic. The Journal of Immunology 190:5306–5312.

Zhang, X., D. Zhao, X. Xiong, Z. He, and H. Li. 2016. Multifaceted Histone H3 Methylation and Phosphorylation Readout by the Plant Homeodomain Finger of Human Nuclear Antigen Sp100C. Journal of Biological Chemistry 291:12786–12798.

Zhou, L., Y. Wang, M. Zhou, Y. Zhang, P. Wang, X. Li, J. Yang, H. Wang, and Z. Ding. 2018. HOXA9 inhibits HIF-1α-mediated glycolysis through interacting with CRIP2 to repress cutaneous squamous cell carcinoma development. Nature Communications. doi:10.1038/s41467-018-03914-5.

Zucchelli, C., S. Tamburri, G. Filosa, M. Ghitti, G. Quilici, A. Bachi, and G. Musco. 2019. Sp140 is a multi-SUMO-1 target and its PHD finger promotes SUMOylation of the adjacent Bromodomain. Biochim Biophys Acta Gen Subj 1863:456–465.

