## Supplemental Figures for "Immune chromatin reader SP140 regulates microbiota and risk for inflammatory bowel disease"

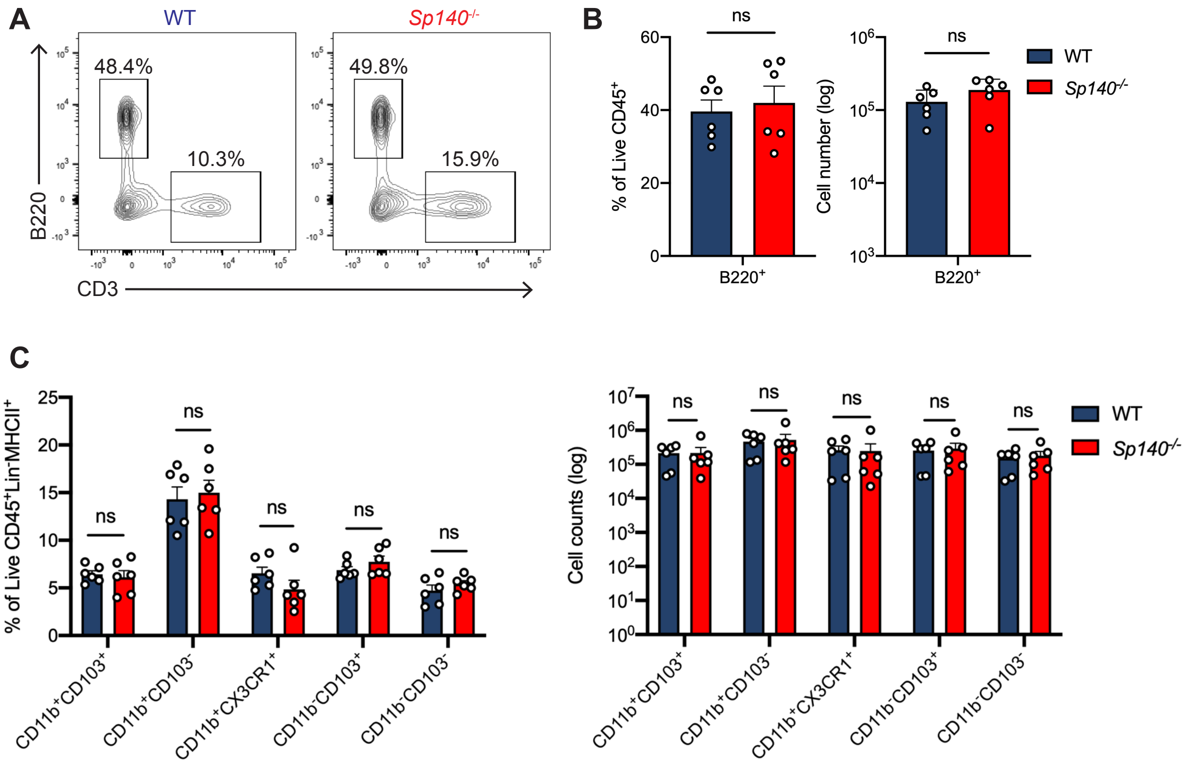


**Figure S1. Characterization of leukocyte populations in WT versus *Sp140*$^{-/-}$ mouse colon lamina propria at homeostasis. (A)** Representative flow cytometry plots of live CD45$^{+}$ cells gated on B220+ and CD3+ populations. **(B)** Quantification of frequency and total count of B220$^{+}$ B cells. **(C)** Quantification of frequency and total count of mononuclear phagocytes. **(A-C)** n=6, *P<0.05, **P<0.01, ***P<0.001, ****P<0.0001; unpaired *t* test. Error bars represent s.e.m.

| Diagnosis | PUB ID | Sex | Age at Diagnosis | Age at Collection | In Remission? | Medication at Time of Collection |
|---|---|---|---|---|---|---|
| Healthy | 160826 | F | N/A | 46 | N/A | N/A |
| Control | 120777 | M | N/A | 22 | N/A | N/A |
| *SP140* $^{wt}$ | 107560 | F | N/A | 22 | N/A | N/A |
| Crohn's | 109307 | F | 10 | 23 | Yes | Remicade |
| Disease | 110577 | F | 38 | 38 | No | Mesalamine |
| *SP140* $^{wt}$ | 138741 | M | 21 | 43 | Yes | Ciprofloxacin, Budesonide, Mercaptopurine, Humira |
| Crohn's | 159126 | F | 46 | 60 | No | Prednisone, Simponi |
| Disease | 184102 | F | 61 | 66 | Yes | Prednisone |
| *SP140* $^{mut}$ | 160916 | F | 15 | 37 | Yes | Cimzia |

**Figure S2. Characterization of healthy control and Crohn's disease (CD) cohort at time of stool collection. (A)** Patient information, including age, remission status, and medication at time of stool collection for healthy control, CD patients expressing wildtype SP140 (SP140$^{wt}$), and CD patients bearing the common SP140 genetic variant (SP140$^{mut}$) (n=3).
